# One Health in practice: Multi-season survey of ixodid tick species collected from domestic dogs in Chad

**DOI:** 10.1101/2024.06.05.597634

**Authors:** Christopher A. Cleveland, Morgan Friedman, Alec T. Thompson, Ellen Haynes, Sarah M. Coker, John A. Bryan, Metinou K. Sidouin, Philip Tchindebet Ouakou, Bongo Nare Richard Ngandolo, Michael J. Yabsley

## Abstract

Ticks are medically important vectors of pathogens, many of which are zoonotic or impact domestic animal and/or wildlife health. Climate change, landscape use, and increasing interactions between domestic animals, wildlife, and humans have resulted in changes in tick-host dynamics and the emergence of novel pathogens worldwide. Therefore, it is crucial to describe the host and geographic ranges of vector species to assess disease risk, particularly in areas where research is lacking. This task necessitates adopting a One Health approach. In sub-Saharan Africa, previous work on ticks has focused primarily on those species most relevant to domestic livestock or humans, highlighting a significant knowledge gap concerning species of ticks that infest peri-domestic animals in rural areas. The objective of this study was to investigate the species diversity of ticks on peri-domestic dogs in rural areas of Chad, Africa. From 2019-2022, we collected 3,412 ixodid ticks from 435 peri-domestic dogs from 23 villages in Chad, Africa during both dry and wet seasons. Ticks were identified to species using morphological techniques and/or molecular analyses of the 16S rRNA, 12S rRNA, and cytochrome oxidase I gene regions. We identified 11 species of ticks from dogs including *Amblyomma variegatum*, *Amblyomma marmoreum*, *Haemaphysalis leachi,* a *Haemaphysalis* sp., *Hyalomma impressum, Hyalomma truncatum, Rhipicephalus decoloratus, Rhipicephalus guilhoni, Rhipicephalus muhsamae, Rhipicephalus linnaei* (=*R. sanguineus* ‘tropical lineage’), and a *Rhipicephalus* sp. Several of these tick species are known vectors for important canine and zoonotic pathogens and some are more commonly associated with cattle hosts. Our results show that sampling ticks from peri-domestic dogs provides an opportunity to examine vectors that may infest domestic animals, agricultural animals, wildlife, and humans as hosts in an understudied area.

**Highlights:**

- Tick-borne diseases are significant concern across sub-Saharan Africa
- Limited data on ticks of animals and people in Chad, Africa
- A high diversity of tick species infested domestic dogs in Chad
- Many detected tick species associated with important pathogens

## Introduction

In response to global climate change, land-use alterations, and globalization of animal trade and human movements, the species diversity and geographical ranges of vectors are changing (Altizer et al., 2013). Specifically, tick ranges are predicted to expand and shift into novel regions, potentially altering host-parasite community dynamics and introducing infectious agents to new hosts and environments (Leger et al., 2013). Ticks are important vectors of viral, bacterial, and protozoan pathogens that can cause severe disease in humans and domestic animals, resulting in predicted annual economic losses upwards of $20 billion USD (Lew-Tabor and Rodriguez Valle, 2016). Additionally, individual tick species can transmit multiple pathogens, leading to coinfections in definitive hosts (Jongejan and Uilenberg, 2004; Walker et al., 2005; Yessinou et al., 2022; Diarra et al., 2023). There are many tick genera in Africa that transmit numerous high-impact pathogens for humans, domestic animals, and wildlife, including the causative agents of ehrlichiosis, anaplasmosis, rickettsioses, theileriosis, babesiosis, and heartwater (Springer et al., 2020).

The control of ticks and prevention of tick-borne diseases require an interdisciplinary and collaborative approach because of the complex natural history of ticks that involves humans, animals, and the environment. This One Health approach has been recognized and adopted by a quadripartite initiative in December 2023 by the World Health Organization (WHO), the Food and Agriculture Organization of the United Nations (FAO), the United Nations Environment Programme (UNEP), and the World Organization for Animal Health (WOAH, formally the Office International des Epizooties (OIE)). Peri-domestic dogs in Chad live at the intersection of various ecological niches, often interacting with humans, agricultural animals, and wildlife, and may be infected with a suite of important tick-borne pathogens. Therefore, these dogs present a unique opportunity to investigate tick ecology in rural, understudied areas of sub-Saharan Central Africa. The goal of this study was to assess the diversity and prevalence of tick species found on peri-domestic dogs in Chad, and to examine associations between tick infestations and spatial and seasonal factors.

## Materials and methods

Ticks were collected from domestic dogs (n = 435) during physical exams as part of a study on Guinea worm (*Dracunculus medinensis*) in Chad, Africa (Cleveland et al., 2022). Collections occurred in May 2019, November 2019, and May 2020 in 23 villages located along the Chari River (Fig. 1). The same individual dogs were resampled at each collection point, except for dogs who were not available due to mortality or movement outside of the original village. Ticks were preserved in 70% ethanol and most were identified to species level according to morphological keys (Hoogstraal, 1956; Estrada-Peña et al., 2004). When necessary, molecular analyses (PCR) were used to confirm some identifications of specimens that were damaged and lacked key morphological characteristics or to identify some immature specimens that could not identified to species due to a lack of comprehensive morphological keys (Table 1). In addition, *Rhipicephalus muhsamae* is part of the *R. simus* species complex and shares many morphological traits with the other species of this group; therefore, molecular analysis was conducted for confirmation of our morphological identification of this species.

**Figure 1.**
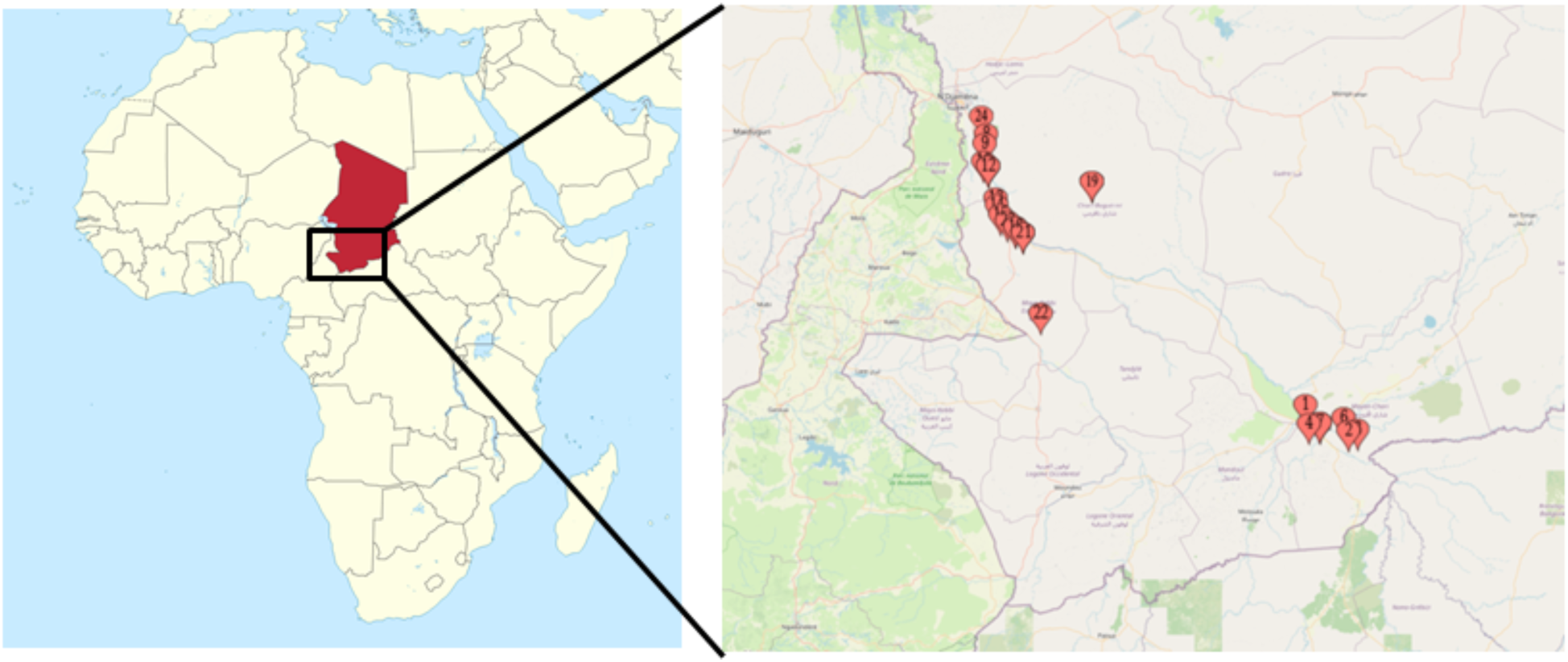
Map of 23 villages in Chad, Africa where ticks were sampled from peri-domestic dogs in 2019 and 2020.

**Table 1.**
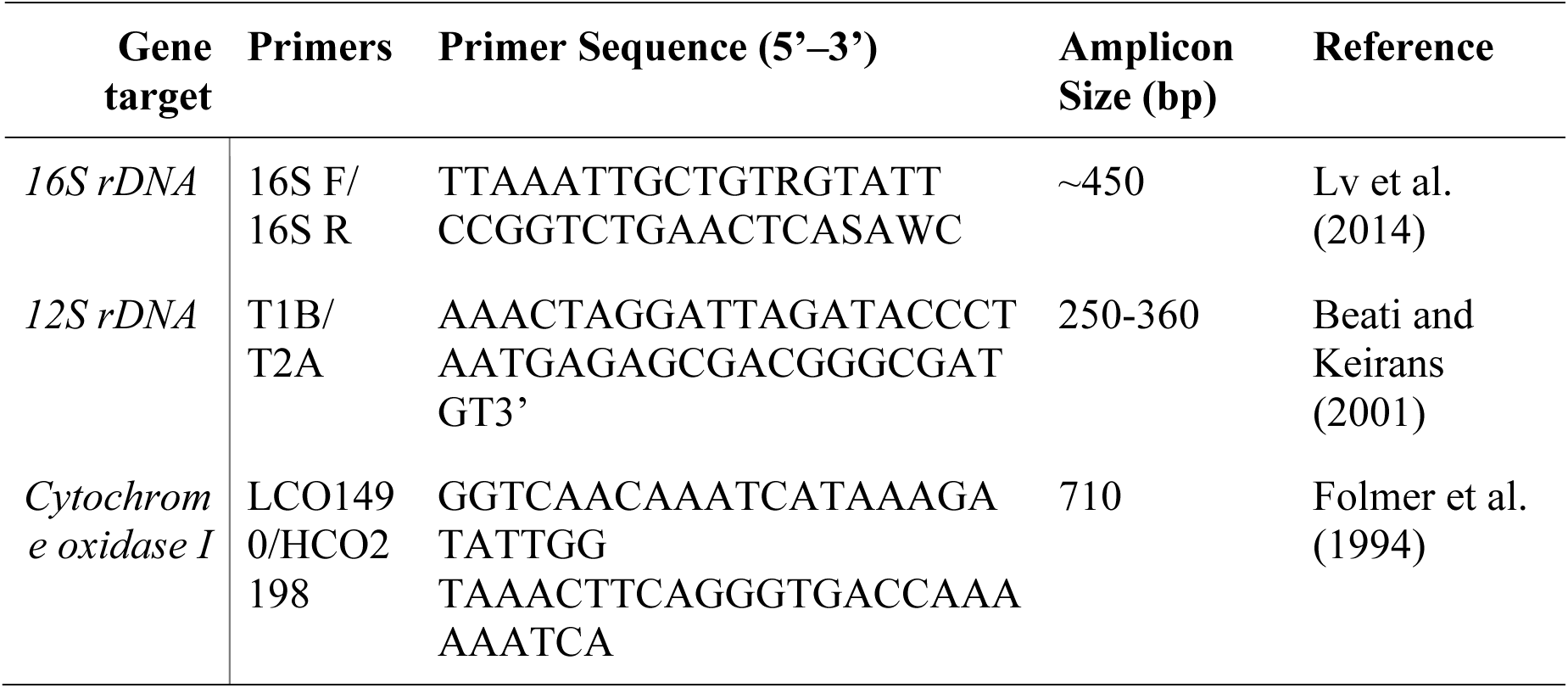
PCR primers used for molecular species-level identification of ticks collected from peri-domestic dogs in Chad, Africa.

For PCR testing, ticks were medially bisected with a sterile razor blade and one half was placed in a dry, open microcentrifuge tube in a biosafety hood at room temperature for 12 hrs to allow residual ethanol to evaporate. The remaining half of each tick was stored in 70% ethanol for future reference. DNA was extracted from the ticks using a DNeasy blood and tissue kit (Qiagen, Hilden, Germany) following the manufacturer’s protocol. DNA was stored at −20 °C until PCR amplification. The 16S rDNA region was the primary gene target for tick species identification. The 12S rDNA gene or cytochrome oxidase I gene targets were used if the 16S PCR failed to amplify or to provide an additional gene region for species confirmation. The tick DNA fragments were amplified using primer sets and conditions previously described (Table 1). Amplicons were visualized on 2% agarose gels stained with GelRed Nucleic Acid Gel Stain (Biotium, Hayward, CA, USA). Positive amplicons were extracted from the gels and purified using the QIAquick gel extraction kit (Qiagen) then submitted for bidirectional Sanger sequencing (GENEWIZ Corporation, South Plainfield, NJ, USA). Chromatographs were analyzed using Geneious Prime version 11.0.12 (Geneious, Auckland, New Zealand, https://www.geneious.com/). Sequences were compiled and compared to preexisting records using the BLASTN program in GenBank (National Center for Biotechnology Information, Bethesda, MD, USA).

Novel *Rhipicephalus* sequences were aligned with related sequences from GenBank using ClustalW and phylogenetic trees were constructed via a maximum likelihood algorithm using MEGA X (Kumar et al., 2018; Stecher et al., 2020) and the Tamura-Nei method (Tamura and Nei, 1993) (Fig. 2). Due to some known inconsistency of *Rhipicephalus* species identification, inaccurate identities are likely present for some individuals in GenBank; however, these sequences were included in our phylogenetic analysis and labeled as noted in GenBank.

**Figure 2.**
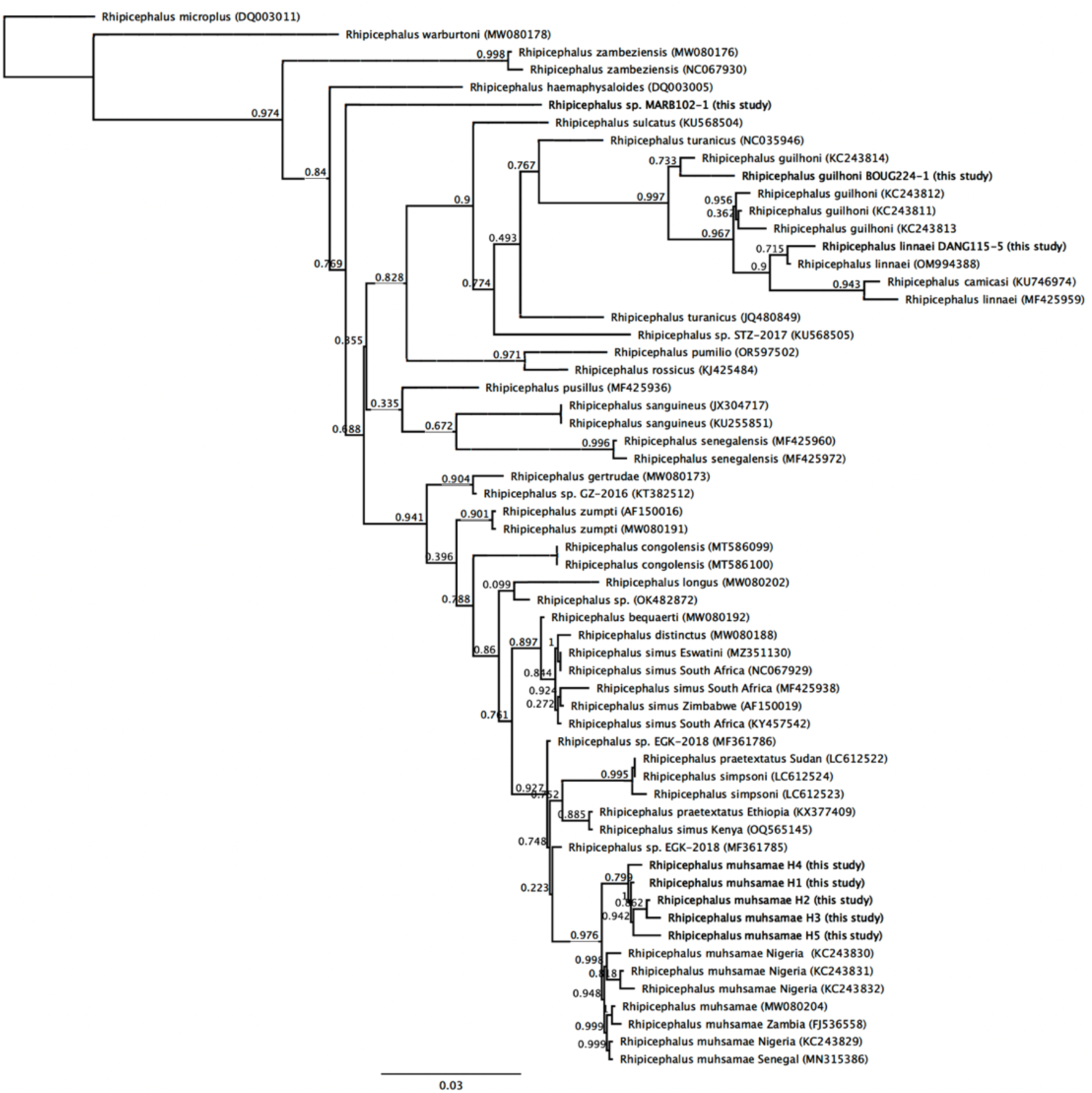
Phylogenetic analysis of 12S rRNA gene sequences for *Rhipicephalus* spp. collected from peri-domestic dogs in Chad, Africa in 2019 and 2020.

Prevalence was calculated as the percentage of dogs with one or more ticks present out of the total number of dogs examined. For each prevalence estimate, 95% confidence intervals were calculated using commercial software.

## Results

Ticks were found on 84% of dogs sampled (Table 2). A total of 3,412 ticks from four genera and 11 species were collected from 2019 to 2020. The ticks included 3,214 adults and 198 nymphs; no larvae were collected. The numbers of ticks of each species collected by time point of sampling, tick life stage, and tick sex are shown in Table 3.

**Table 2.** Prevalence of ticks on peri-domestic dogs in Chad, Africa at three time points in 2019 and 2020. Note that some dogs had multiple tick species present. (n = number of Chadian dogs sampled in each collection period)

| Species | May 2019<br>(n=435) | November 2019<br>(n=318) | May 2020<br>(n=254) | Total<br>(n=1007) |
| --- | --- | --- | --- | --- |
| *Amblyomma marmoreum* | 2<br>(<1%) | 6<br>(1.9%) | 1<br>(<1%) | 9<br>(1%) |
| *Amblyomma variegatum* | 11<br>(2.5%) | 60<br>(18.9%) | 6<br>(2.4%) | 77<br>(7.6%) |
| *Haemaphysalis leachi* | 4<br>(<1%) | 7<br>(2.2%) | 3<br>(1.2%) | 14<br>(1.4%) |
| *Haemaphysalis* sp. | - | 1<br>(<1%) | - | 1<br>(<1%) |
| *Hyalomma impressum* | - | 1<br>(<1%) | - | 1<br>(<1%) |
| *Hyalomma truncatum* | - | 3<br>(1%) | 1<br>(<1%) | 4<br>(<1%) |
| *Rhipicephalus decoloratus* | - | 2<br>(<1%) | - | 2<br>(<1%) |
| *Rhipicephalus guilhoni* | - | - | 2<br>(<1%) | 2<br>(<1%) |
| *Rhipicephalus muhsamae* | 34<br>(7.8%) | 164<br>(51.6%) | 55<br>(21.6%) | 253<br>(25.1%) |
| *Rhipicephalus linnaei* | 287<br>(66.0%) | 257<br>(80.8%) | 227<br>(89.4%) | 771<br>(76.6%) |
| *Rhipicephalus* sp. | 2<br>(<1%) | - | 1<br>(<1%) | 3<br>(<1%) |
| Total number of dogs with ticks | 300<br>(69.0%) | 302<br>(95.0%) | 241<br>(94.9%) | 843<br>(83.7%) |

**Table 3.**
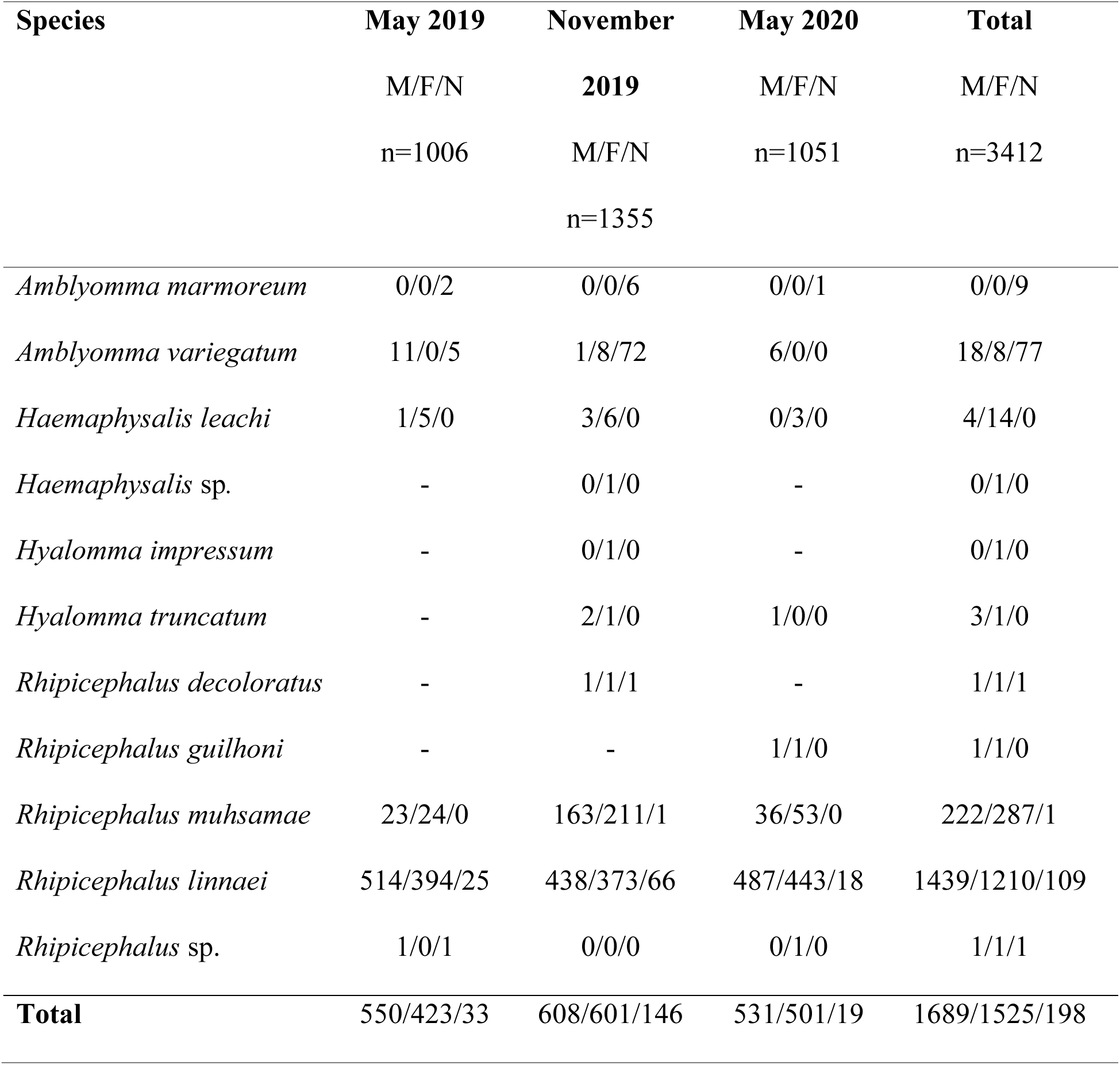
Species and life stages of ticks collected from peri-domestic dogs in Chad, Africa in 2019 and 2020. Results are shown by time point of collection, the total number collected for the entirety of the study, and further divided by tick sex/life stage. M = Adult male, F = Adult female, N = Nymph, n = total number of ticks collected, dashes indicate no ticks of that species collected during that time point.

The most frequently encountered tick species on dogs was *R. sanguineus* sensu lato, which were present on 77% of dogs during all three time periods. They comprised the majority of ticks collected, accounting for 2,758 out of 3,412 ticks (80.8%) (Tables 2 and 3). Sequence analysis of *R. linnaei* (formerly tropical lineage of *R. sanguineus)* (Fig. 2 and 3) (Slapeta et al., 2021; Slapeta et al., 2022). The next most common tick collected was *R. muhsamae* (25.1% of dogs infested and 14.9% of ticks collected). A relatively high percentage (7.6%) of dogs were infested with low numbers of *A. variegatum*. The remaining tick species were all found at very low prevalences (<1.5%) (Table 2).

**Figure 3.**
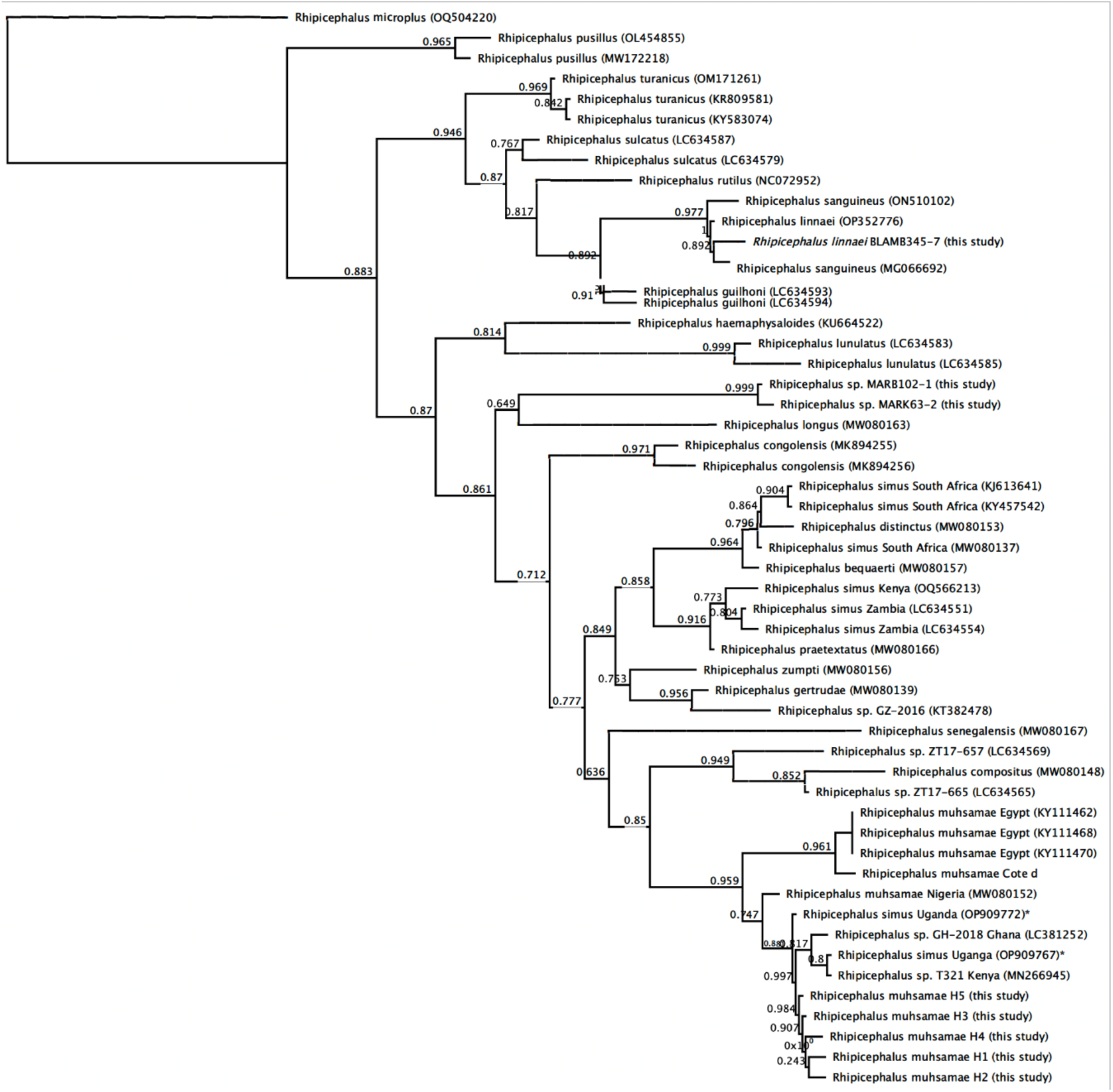
Phylogenetic analysis of 16S rRNA gene sequences for *Rhipicephalus* spp. collected from peri-domestic dogs in Chad in 2019 and 2020. Two sequences of *R. simus* (marked with *) are considered misidentifications based on data from Walker et al. (2005) who state *R. simus* s.s. are restricted to southern Africa.

The second most common tick was morphologically and genetically identified as *R. muhsamae*, a species that is part of the *R. simus* species complex. There were five unique 16S rRNA gene haplotypes identified among these specimens that each differed from each other by 1 bp (381/382 bp, 99.7% similar). These sequences were most similar (98.7-100%) to numerous *R. muhsamae* sequences in GenBank, as well as some erroneously submitted as *R. simus* based on the data from Walker et al. (2005) that states *R. simus* sensu stricto are restricted to southern Africa (marked with * in Figure 3). Our sequences were only 93.5-94.5% similar to several *R. muhsamae* sequences from Egypt (KY111455-KY111471). Phylogenetic analysis showed that our sequences grouped together and in the same clade with *R. muhsamae* (Figure 3). These analyses confirmed that samples of *R. simus* from southern Africa grouped separately from *R. muhsamae* and “*R. simus*" samples from West and Central Africa (i.e., Cote d’Ivoire, Egypt, Ghana, Kenya, Nigeria, and Uganda) (Figure 3). Sequences of 12S rRNA also revealed five unique haplotypes that were 99.4-99.7% similar to each other and 98-99.4% similar to various *R. muhsamae* sequences in GenBank. Phylogenetically, these sequences grouped together and as a sister clade to Central African populations of *R. muhsamae* (Figure 2). Two individual *Rhipicephalus* sp. nymphs were genetically classified because an identification could not be made morphologically. The 401bp 16S sequences were 99.8% similar to each other. These sequences were most similar (∼92.5-94%) to numerous species in the *R. sanguineus* s.l. group and, phylogenetically, they grouped together along with *R. longus* (Figure 3). The 12S rRNA gene sequence from one tick was most similar to several members of the *R. sanguineu*s s.l. group (93-95.3%). Phylogenetically, it was on a basal branch to the *R. sanguineus* s.l. group so the species of this tick is unknown (Figure 2).

Eight nymphal *Amblyomma* ticks were identified as *A. marmoreum* by analysis of the 12S sequences (348 bp). Seven of these sequences were identical and the remaining one sequence was 99.1% similar to the other sequences. Our sequences were 98.8-99.4% identical to *A. marmoreum* collected from migratory bird species in Italy and Malta (KC817413, OL352895, and OM522963-OM522980) (Toma et al., 2014). Similarly, the 16S rRNA gene (405 bp) was most similar (99.5%) to numerous *A. marmoreum* sequences from ticks from birds in Italy and a *Kinixys* sp. from South Africa (MW290507, OM462845-OM462862) (Mofokeng et al., 2021). Phylogenetically, our *A. marmoreum* sequences for both gene targets grouped with other *A. marmoreum* sequences (Figures 4 and 5).

**Figure 4.**
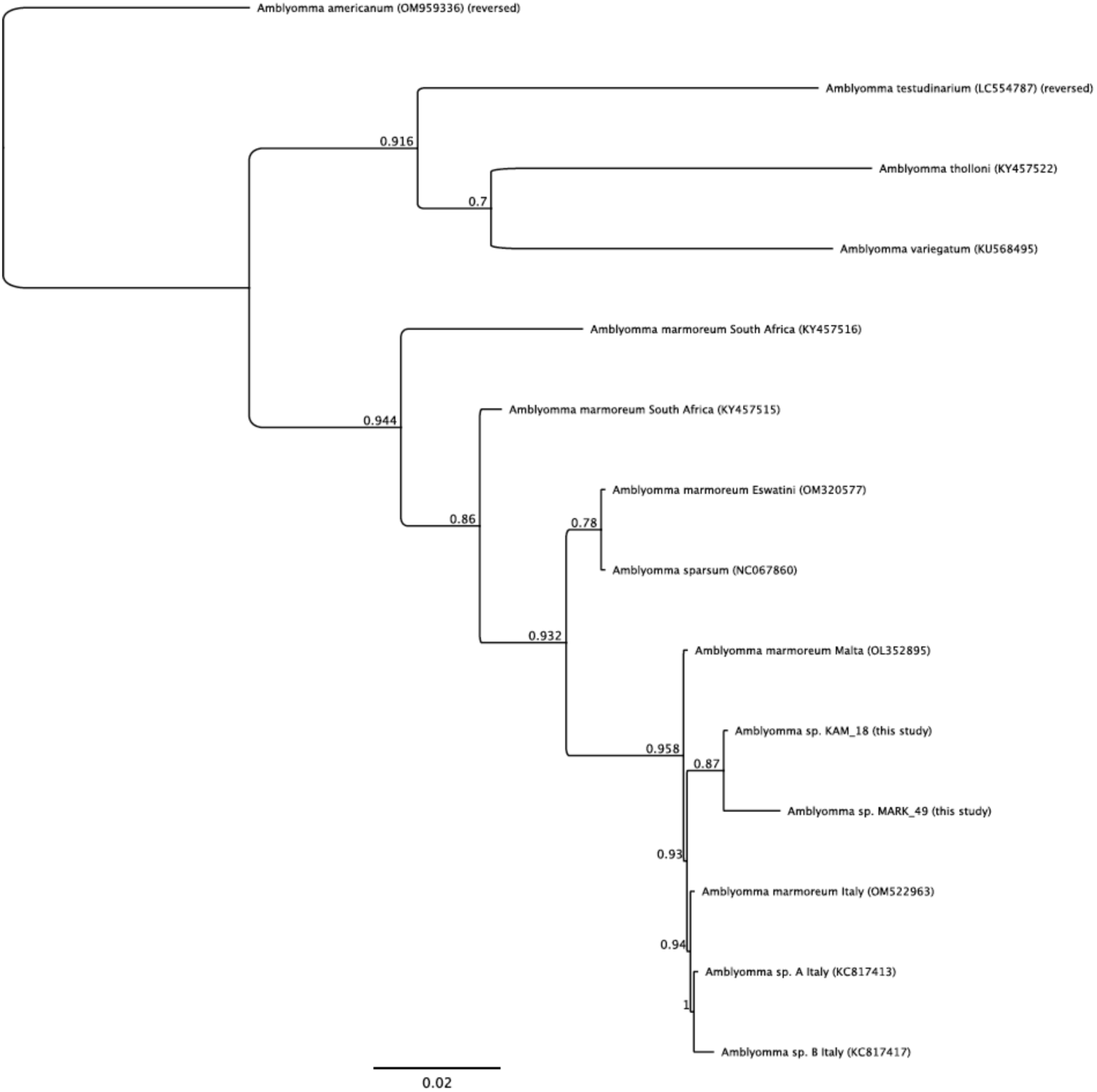
Phylogenetic analysis of 12S rRNA gene sequences for *Amblyomma marmoreum* and related *Amblyomma* spp. collected from peri-domestic dogs in Chad in 2019 and 2020.

**Figure 5.**
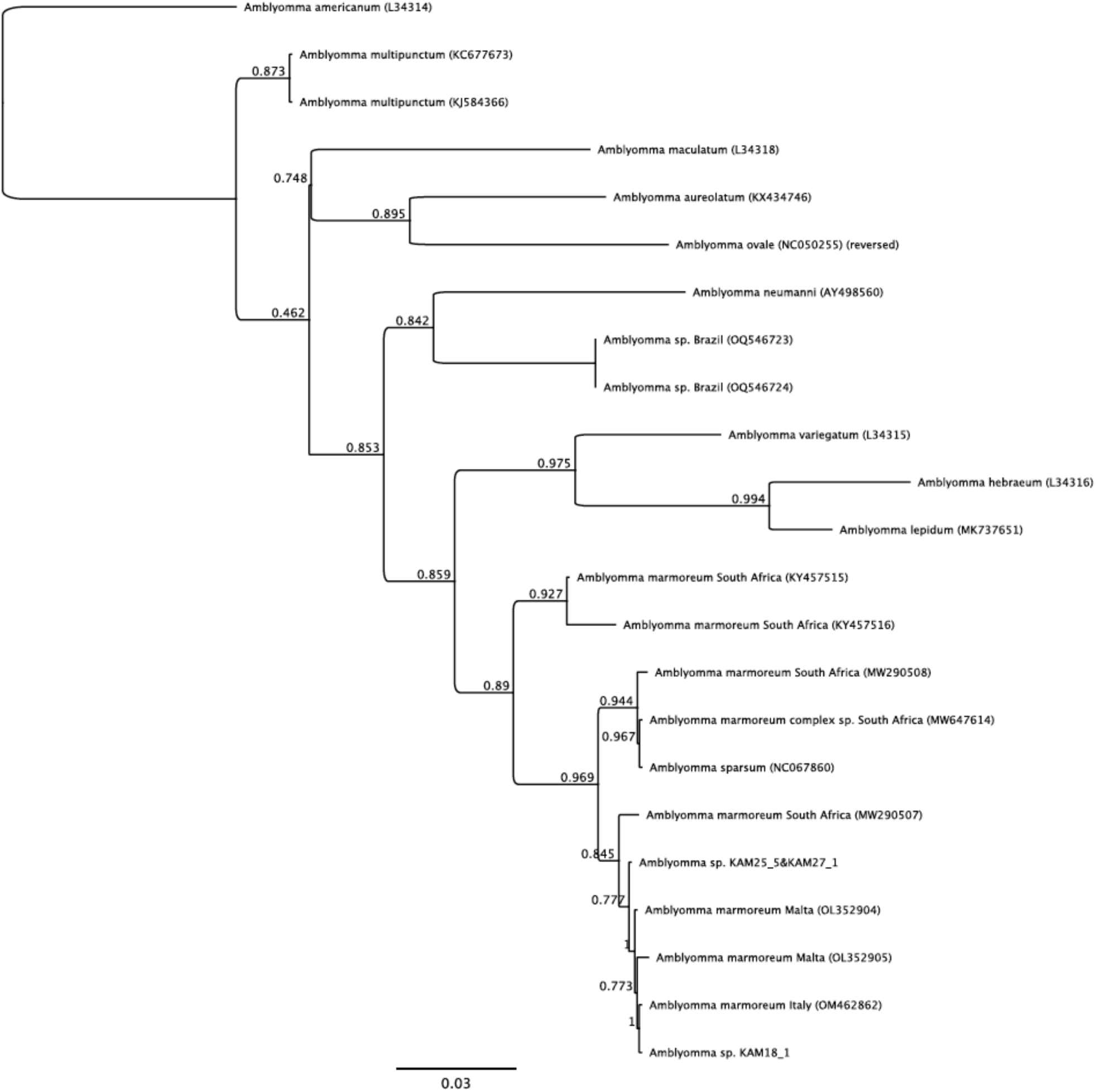
Phylogenetic analysis of 16S rRNA gene sequences for *Amblyomma marmoreum* and related *Amblyomma* spp. collected from peri-domestic dogs in Chad in 2019 and 2020.

A single adult female tick that was markedly damaged (only ¼ of the posterior end collected) was identified as a *Haemaphysalis* sp. by molecular characterization. The 16S rRNA gene sequence (400 bp) was most similar to several *Haemaphysalis* species from rodents in Saudi Arabia [e.g., MW763058, (Alghamdi et al., 2021)] followed by *H. hispanica* (355/380bp, 93.4%) from rabbits in Portugal (OL656102) and Spain (MZ420715) (Figure 6). The partial cytochrome oxidase I gene sequence (709 bp) for this *Haemaphysalis* sp. was most similar to *H. concinna* (569/649 bp, 87.7%) and *H. bispinosa* (599/687 bp, 87.2%).

**Figure 6.**
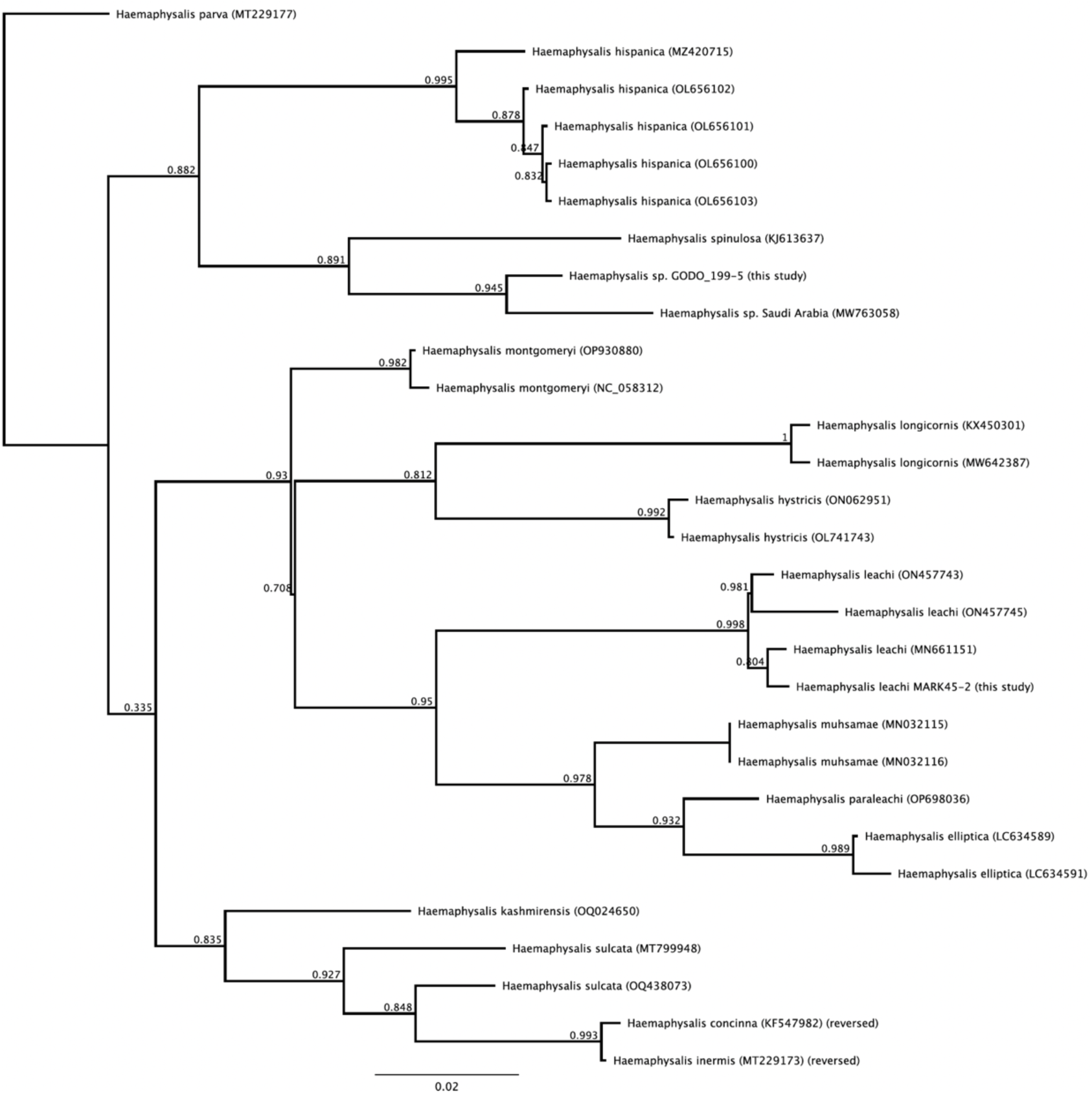
Phylogenetic analysis of 16S rRNA gene sequences for *Haemaphysalis* spp. collected from peri-domestic dogs in Chad in 2019 and 2020.

The overall species diversity of ticks sampled was greatest in November 2019, with seven unique species identified from four genera. There was a higher percentage of examined dogs infested with ticks in November 2019 (95.0%) and May 2020 (94.9%) compared to May 2019 (69.0%) (Table 3). This trend was true for both the northern region and southern regions (Table 4, Figure 7). For *R. muhsamae* specifically, there was a higher percentage of dogs with this tick in the northern region, compared to the southern region, at every time point of the study (Table 4, Figure 4). Variable prevalence was also noted between villages; however, low sample sizes in each village precluded robust statistical analysis (Table 5).

**Figure 7.**
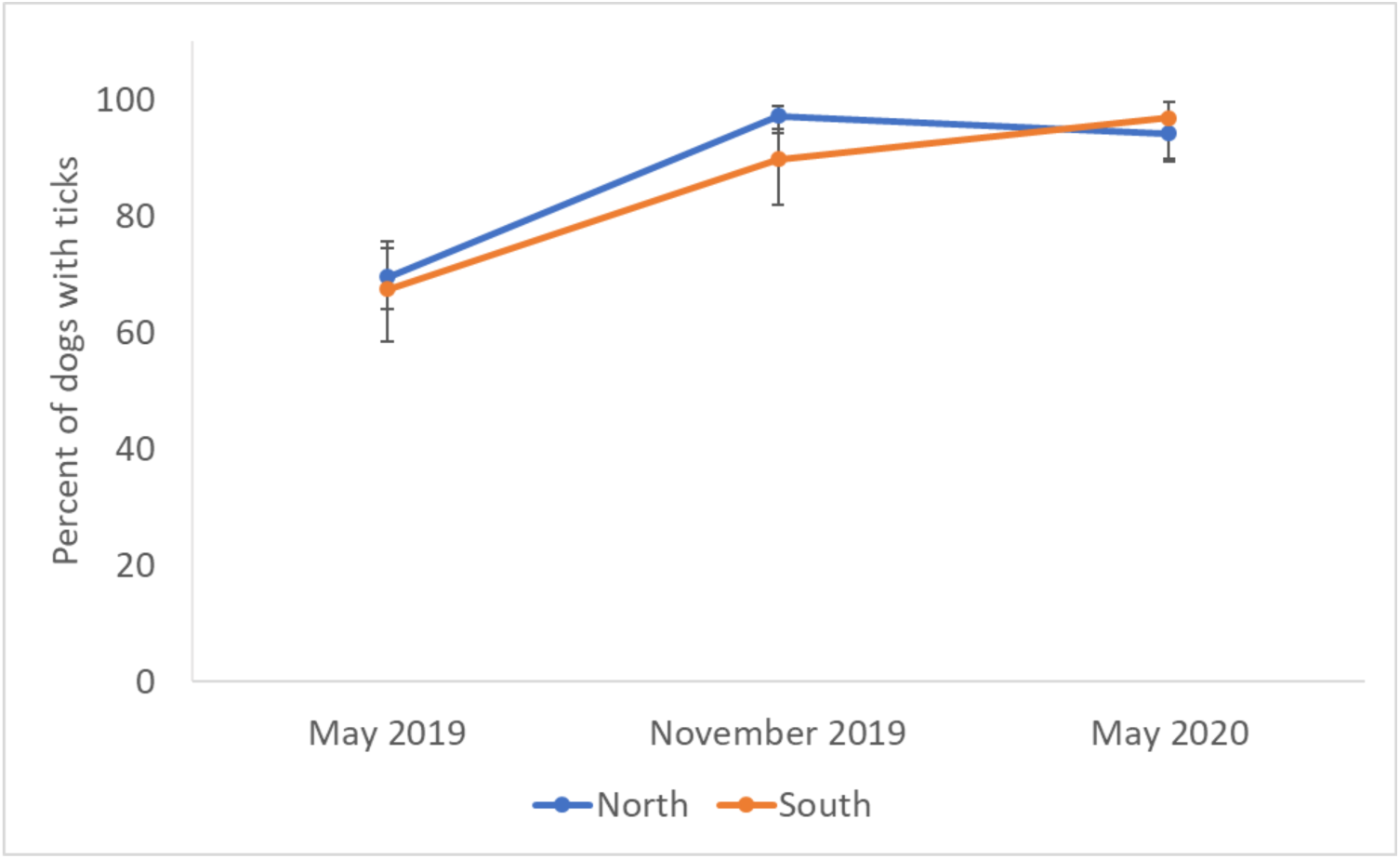
Percentage of peri-domestic dogs in the north and south regions with ticks present at each of the three timepoints of the study. Error bars show 95% confidence intervals for percent prevalence.

**Figure 8.**
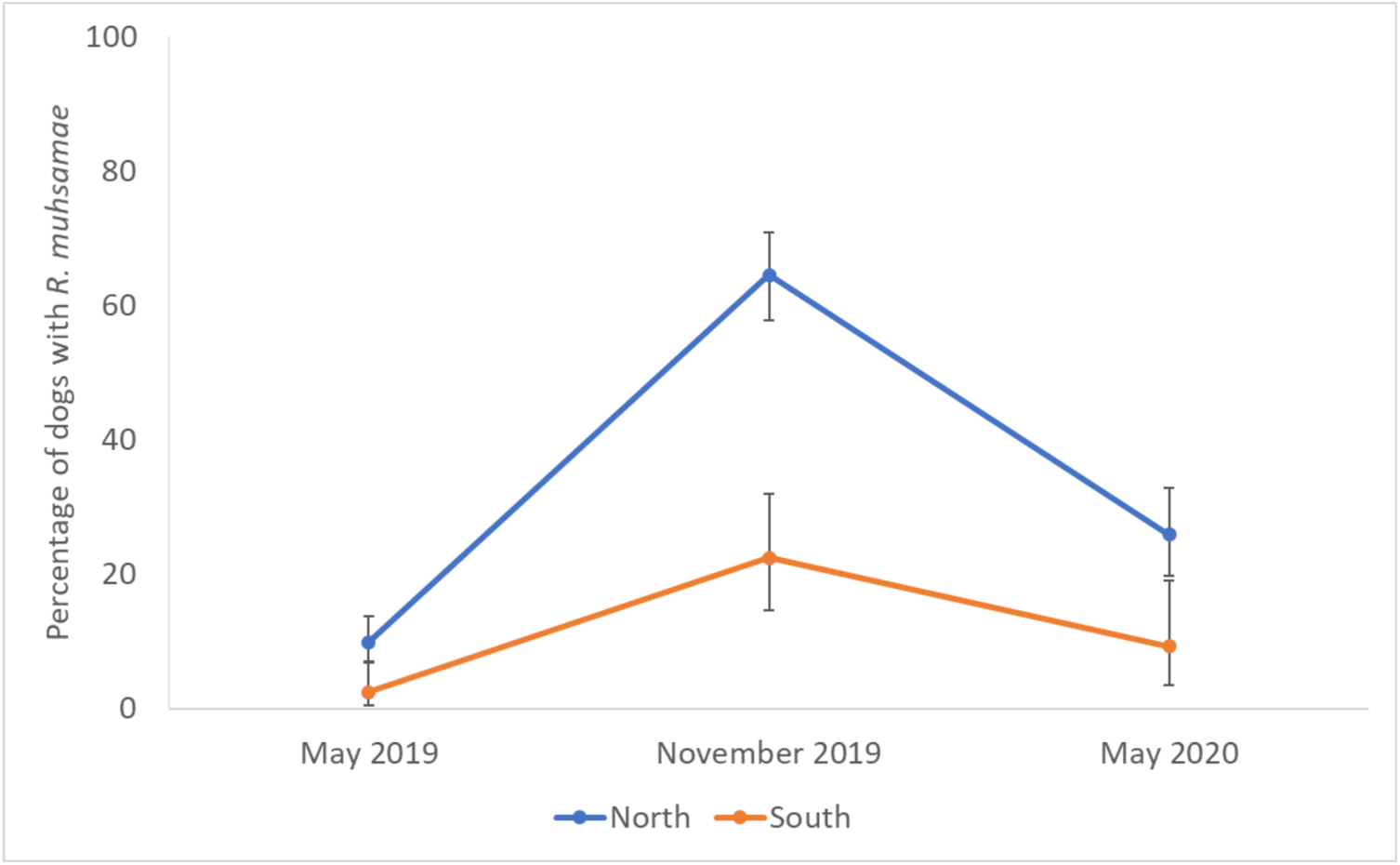
Percentage of peri-domestic dogs in the north and south regions of Chad with *R. muhsamae* present at each of the three timepoints of the study. Error bars show 95% confidence intervals for percent prevalence.

**Table 4.** Prevalence of ticks (overall and for *Rhipicephalis muhsamae*) on peri-domestic dogs in Chad, Africa at three time points in 2019-2020 by region. 95% confidence intervals are provided.

|  | May 2019 | November 2019 | May 2020 |
| --- | --- | --- | --- |
| **All ticks** |  |  |  |
| North region | 69.6% (64.1-74.6) | 97.3% (94.2-99.0) | 94.2% (89.8-97.1) |
| South region | 67.5% (58.4-75.6) | 89.8% (82.0-95.0) | 96.9% (89.3-99.6) |
| ***R. muhsamae*** |  |  |  |
| North region | 9.9% (6.8-13.8) | 64.5% (57.8-70.9) | 25.9% (19.8-32.8) |
| South region | 2.4% (0.5-7.0) | 22.4% (14.6-32.0) | 9.2% (3.5-19.0) |

**Table 5.** Prevalence of ticks (overall and for *R. linnaei, R. muhsamae, and A. variegatum*) on peri-domestic dogs in Chad, Africa across the 23 villages sampled. 95% confidence intervals are provided in parentheses.

| Village Name | Number of dogs sampled | Prevalence of any ticks | Prevalence of *Rhipicephalus linnaei* | Prevalence of *Rhipicephalus muhsamae* | Prevalence of *Amblyomma variegatum* |
| --- | --- | --- | --- | --- | --- |
| **North Region** |  |  |  |  |  |
| 5Kilos | 16 | 93.8%<br>(69.8-99.8) | 56.3%<br>(29.9-80.2) | 75.0%<br>(47.6-92.7) | 50.0%<br>(24.6-75.4) |
| Abechere | 4 | 100.0%<br>(39.8-100.0) | 100.0%<br>(39.8-100.0) | 75.0%<br>(19.4-99.4) | 75.0%<br>(19.4-99.4) |
| Abory Mbarma | 9 | 66.7%<br>(29.9-92.5) | 55.6%<br>(21.2-86.3) | 33.3%<br>(7.5-70.1) | 11.1%<br>(0.3-48.2) |
| Aligarga Sara | 22 | 100.0%<br>(84.6-100.0) | 100.0%<br>(84.6-100.0) | 22.7%<br>(7.8-45.4) | 0.0%<br>(0.0-15.4) |
| Boudanassa Bao | 16 | 81.3%<br>(54.4-96.0) | 81.3%<br>(54.4-96.0) | 43.8%<br>(19.8-70.1) | 18.8%<br>(4.0-45.6) |
| Boudanassa Lambe | 38 | 100.0%<br>(90.8-100.0) | 100.0%<br>(90.8-100.0) | 68.4%<br>(51.4-82.5) | 15.8%<br>(6.0-31.2) |
| Bougemene | 24 | 95.8%<br>(78.9-99.9) | 95.8%<br>(78.9-99.9) | 75.0%<br>(53.3-90.2) | 4.2%<br>(0.1-21.1) |
| Darda | 20 | 90.0%<br>(68.3-98.8) | 90.0%<br>(68.3-98.8) | 50.0%<br>(27.2-72.8) | 10.0%<br>(1.2-31.7) |
| Godogo | 15 | 100.0%<br>(78.2-100.0) | 100.0%<br>(78.2-100.0) | 60.0%<br>(32.3-86.7) | 0.0%<br>(0.0-21.8) |
| Kakalembéri | 27 | 96.3%<br>(81.3-99.9) | 96.3%<br>(81.3-99.9) | 63.0%<br>(42.4-80.6) | 18.5%<br>(6.3-38.1) |
| Magrao | 15 | 86.7%<br>(59.5-98.3) | 73.3%<br>(44.9-92.2) | 86.7%<br>(59.5-98.3) | 6.7%<br>(0.2-32.0) |
| Marabe 2 | 31 | 83.9%<br>(66.3-94.6) | 77.4%<br>(58.9-90.4) | 19.4%<br>(7.4-37.5) | 19.4%<br>(7.4-37.5) |
| Mecontie | 20 | 90.0%<br>(68.3-98.8) | 85.0%<br>(62.1-96.8) | 40.0%<br>(19.1-64.0) | 30.0%<br>(11.9-54.3) |
| Mogrom Biao | 11 | 90.9%<br>(58.7-99.8) | 90.9%<br>(58.7-99.8) | 63.6%<br>(30.8-89.1) | 9.1%<br>(0.2-41.3) |
| Mogrom Centre | 10 | 90.0%<br>(55.5-99.8) | 90.0%<br>(55.5-99.8) | 70.0%<br>(34.8-93.3) | 10.0%<br>(0.2-44.5) |
| Raf Centre | 15 | 100.0%<br>(78.2-100.0) | 100.0%<br>(78.2-100.0) | 60.0%<br>(32.3-86.7) | 0.0%<br>(0.0-21.8) |
| Sawata Matoum | 19 | 89.5%<br>(66.9-98.7) | 68.4%<br>(43.4-87.4) | 73.7%<br>(48.8-90.8) | 31.6%<br>(12.6-56.6) |
| **South Region** |  |  |  |  |  |
| Bodobo | 36 | 97.2%<br>(85.5-99.9) | 97.2%<br>(85.5-99.9) | 0.0%<br>(0.0-9.7) | 27.8%<br>(14.2-45.2) |
| Dangala Kanya | 16 | 93.8%<br>(69.8-99.8) | 93.8%<br>(69.8-99.8) | 25.0%<br>(7.3-52.4) | 18.8%<br>(4.0-45.6) |
| Dankolo | 12 | 100.0%<br>(73.5-100.0) | 100.0%<br>(73.5-100.0) | 0.0%<br>(0.0-26.5) | 16.7%<br>(2.1-48.4) |
| Doumabe | 12 | 58.3%<br>(27.7-84.8) | 58.3%<br>(27.7-84.8) | 41.7%<br>(15.2-72.3) | 33.3%<br>(9.9-65.1) |
| Kaimambe | 18 | 94.4%<br>(72.7-99.9) | 94.4%<br>(72.7-99.9) | 27.8%<br>(9.7-53.5) | 11.1%<br>(1.4-34.7) |
| Marakouya 2 | 29 | 100.0%<br>(88.1-100.0) | 96.6%<br>(82.2-99.9) | 41.4%<br>(23.5-61.1) | 6.9%<br>(0.8-22.8) |

## Discussion

This study provides new information on the prevalence and diversity of tick species on domestic dogs in Chad, an area with limited data on ticks and tick-borne diseases. Studying ticks within a One Health framework is crucial because tick-borne diseases affect both humans and animals. Ticks have a complex natural history, involving specific requirements like host species, habitat types, and climatic factors. Of significance to veterinary health, various tick vectors capable of carrying important pathogens were identified, including *R. linnaei, R. decoloratus, A. variegatum, H. leachi*, and *H. truncatum*. Although the latter four ticks were uncommonly detected on dogs, their presence indicates a potential risk to cattle health. In addition, several of these tick species transmit pathogens of consequence to human health. This study indicates that there is a risk for tick-borne diseases in this region of Chad and the noted seasonal and regional differences should be further examined given expected changes in climate, environment, and land use in the region.

A high prevalence and diversity of ticks were detected on peri-domestic dogs in rural areas of Chad, Africa. Although the primary tick species found was *Rh. linnaei* (the brown dog tick), we also found 10 other tick species that commonly infest a broader range of hosts, including humans and agriculturally important species such as cattle. Similar results were found in a recent multi-country study in sub-Saharan Africa in which at least 13 tick species were detected on domestic dogs (Heylen et al., 2021). Our study sites were all rural and Heylen et al. (2021) reported a higher diversity of ticks on dogs from rural sites compared to urban areas; that study also reported that the highest number of ticks were found on dogs from Nigeria, a neighboring country to Chad. The high prevalence and diversity of ticks is important because many of the detected species are important vectors of pathogens. For example, *R. linnaei* transmits several canine pathogens (e.g., *Ehrlichia canis, Anaplasma platys, Babesia* spp., *Hepatozoon* spp.), all of which were detected in dogs we sampled for ticks (Haynes et al., In review). In addition, several of the species collected in this study are vectors of zoonotic and agricultural pathogens (e.g., *Babesia bigemina*, *Anaplasma marginale*, *Ehrlichia ruminantium, Rickettsia africae, R. massiliae*, *R. conorii*, and Crimean-Congo hemorrhagic fever virus (Logan et al., 1989; Parola et al., 2001; Dantas-Torres, 2010). In a concurrent study, several zoonotic *Rickettsia* spp. were detected in ticks collected in this study (Osip et al., In review).

The most common ticks detected on dogs was *R. linneai,* a member of the *R. sanguineus* s. l. group that was historically called the ‘tropical lineage’ (Slapeta et al., 2022). This is not surprising as this is a domestic dog-associated tick species and *R. sanguisuga* s.l. is the most common tick group found on dogs in Africa, including Nigeria where a recent molecular study confirmed that *R. linneaei* was only lineage detected (Kamani, 2021b; Elelu et al., 2022). This species complex has been associated with numerous pathogens of domestic dogs and humans, but given the taxonomic uncertainties of this group, more work is needed to determine the diversity of pathogens transmitted by each species or genotype (Dantas-Torres et al., 2013; Kamani, 2021b; Elelu et al., 2022).

Several cattle associated *Rhipicephalus* spp. (*R. muhsamae, R. guilhoni* and *R. decoloratus*) were detected on dogs. The blue tick, *R. decoloratus*, is a one-host tick that is an important vector of *Babesia bigemina* and *Anaplasma marginale*. It is one of the most abundant cattle ticks in Nigeria (Lorusso et al., 2013). The second most common tick found on dogs in our study, *R. muhsamae*, is also common on cattle in East and Central Africa (Walker et al., 2005).

Although no veterinary pathogens have been associated with this tick species, it is a potential vector of the zoonotic *R. massiliae* (Parola et al., 2013). Only two dogs were infested with *R. guilhoni,* but other studies have found this species on ungulates, various wild carnivores and domestic dogs (Walker et al., 2005; Petney et al., 2017). Although Crimean-Congo hemorrhagic fever virus and *R. massiliae* have been detected in *R. guilhoni*, the vectoral capacity is unknown (Petney et al., 2017).

A low prevalence and intensity of *A. marmoreum* nymphs were detected on dogs in Chad. This tick, the African tortoise tick, is predominantly found on tortoises in southern Africa but has rarely been detected on birds, small mammals, and wild carnivores (Walker, 1991). Sequences from our ticks shared the highest identity with *A. marmoreum* collected from migratory birds in Italy and Malta (Toma et al., 2014; Hornok et al., 2022). A report on a non-migratory species of *A. marmoreum* in Chad would be a large geographic extension of this tick’s range; however, this tick is part of a species complex that also includes *A. falsomarmoreum, A. paulopunctatum, A. nuttalli* and *A. sparsum,* all of which can be difficult to distinguish morphologically, especially the larval and nymphal stages. Although *A. marmoreum* reports on terrestrial hosts are typically from Southern Africa, other members of this complex are in Central Africa (i.e., Central African Republic and Kenya) (Uilenberg et al., 2013; Omondi et al., 2017). Our sequences were more similar to *A. marmoreum* compared to the single sequences available for A*. nuttalli* and *A. sparsum*; however, additional studies combining careful morphological and genetic analysis of ticks in this complex are needed.

A low prevalence and intensity of *Haemaphysalis* and *Hyalomma* spp. were detected on dogs. The yellow dog tick (*H. leachi*), is adapted to domestic dogs as the primary host, similar to *R. sanguineus* s.l.. This tick is predominately found in tropical and subtropical regions of sub-Saharan Africa and has been reported in South Chad and Nigeria (Walker et al., 2005; Kamani et al., 2019). This tick is of importance to dogs as it is a presumed vector of *Babesia rossi* (Kamani, 2021a). A single damaged *Haemaphysalis* sp. could not be identified to species based on the two gene targets analyzed, but it was distinct from *Haemaphysalis* spp. (i.e., *H. leachi, H. paraleachi, H. elliptica, H. spinulosa*) commonly reported from dogs in sub-Saharan Africa. Sequences of two gene targets did not match any expected species either with the closest matches being sequences from ticks from rodents in Saudi Arabia. Therefore, additional surveys of dogs would be needed to find specimens for comprehensive morphologic analysis (Kamani et al., 2019; Heylen et al., 2021). The primary hosts for the two *Hyalomma* spp. (*H. truncatum* and *H. impressum)* are cattle and other large ungulates, but *H. truncatum* has also been reported from domestic dogs (Walker et al., 2005). Both species occur in southern Chad and *H*. *truncatum* is associated with several pathogens of importance to humans and horses and predisposes cattle to myiasis (Walker et al., 2005).

Seasonal variation in tick species diversity was noted. We observed a higher diversity of tick species in November 2019, which is during the cooler and more wet season in Chad. This observation aligns with previous studies that have reported greater tick diversity and frequency during wet seasons in neighboring Nigeria and other parts of Africa (Bayer and Maina, 1984; Zieger et al., 1998; Zeleke and Bekele, 2004; Kamani et al., 2019; Jajere et al., 2023). Weather and general climate trends play a large role in determining the distribution of vector-borne diseases. It is predicted that climate change will lead to changes in both the range and transmission periods of many vector-borne diseases, altering their prevalence as well (Altizer et al., 2013; Caminade et al., 2019). The observed difference in tick diversity between the wet and dry seasons indicates that sub-Saharan tick populations may be particularly responsive to future climate change in the region. We also noted annual differences in prevalence between May 2019 and May 2020, mainly driven by prevalence in *R. linnaei.* The lower prevalence of ticks in May 2019 is unknown but could be due to temperature or rainfall differences or less intense sampling by field personnel during the first round of sampling. Further work is needed to document tick populations over time to determine if there are changes associated with shifting climatic factors.

Most tick species were detected too commonly (*R. linnaei*) or too infrequently (other tick species except for *R. muhsamae*) to investigate spatial patterns. However, for *R. muhsamae* specifically, ticks were more likely to be detected in the north region, compared to the south region, during the first two time points of sampling. Because this is a cattle-associated tick, the presence and density of livestock plays a key role in maintaining this tick in an area. The marked increased prevalence of this tick on dogs in the November sampling period may be related to the peak seasonal activity for adult *R. muhsamae* or increased cattle-dog interactions during this time period (Cornet et al., 1986). A previous study by Zachee et al. (2020) examining tick species on cattle in N’Djamena found five tick species, primarily *Hyalomma marginatum* and *H. lusitanicum,* but fewer numbers of *A. variegatum*. Although *Hyalomma* spp. are the predominate ticks on cattle in Chad in that study, we found very few *Hyalomma* spp. specimens among our sampled dog hosts which supports their preference for cattle hosts. Thus, multi-host surveillance is an ideal way to investigate the tick diversity in an area.

In summary, a high diversity and prevalence of ticks were detected on domestic dogs in Chad, many of which are important vectors for pathogens of public health and veterinary concern. However, this study did have some limitations. This study was done during a concurrent study on the efficacy of flubendazole as a therapeutic for guinea worm infections in domestic dogs (Cleveland et al., 2022). Because of this, village selections and timing of sampling were based on Guinea worm study related factors, so they did not represent all of the important habitat types or seasons that would be of interest in a study on ticks. One concern with this dataset was that ticks were being collected from dogs enrolled in a therapeutic trial. In general, benzimidazoles are not used for ectoparasite control and in support of this, we did not see a decrease in ticks during subsequent sampling periods nor did we see a difference in tick prevalence/diversity between treatment and control dogs. Despite these limitations, important data was gathered on ticks in our sampling regions of Chad. To better understand the risk of tick-borne diseases for people and animals in Chad, future work should focus on sampling a higher diversity of habitats, hosts, and times of year.

## Acknowledgments

As part of concurrent Guinea worm research in Chadian dogs, this work was supported by The Carter Center. A full listing of Carter Center supporters is available at http://www.cartercenter.org/donate/corporate-government-foundation-partners/index.html. Additional support was provided by the wildlife management agencies of the Southeastern Cooperative Wildlife Disease Study member states through the Federal Aid to Wildlife Restoration Act (50 Stat. 917). We thank the team members of Afrique OneASPIRE (http://afriqueoneaspire.org/), Institut de Recherche en Elevage pour le Developpement (IRED), for their outstanding contributions in facilitating this work in the field, as well as the technical staff of The Carter Center and the Programme National d’Eradication du Ver de Guinee, Ministry of Health, N’Djamena, Chad for their support. In particular, we acknowledge Hubert Zirimwabagabo, Mario Romero, and Karmen Unterwegner at The Carter Center for supporting and facilitating this research.

## Notes

### Competing Interest Statement

The authors have declared no competing interest.

